# ARIPA: Agentic Reasoning for Image-based Pathology Analysis

**DOI:** 10.64898/2026.11.16.767159

**Authors:** Qiangqiang Gu, Rong Xia, Rose Doerfler, Jie Chen, Arjun Singh, Sahussapont J. Sirintrapun, Aatur D. Singhi, Ioannis K. Zervantonakis, Brian Isett, Riyue Bao

**Affiliations:** Department of Pathology, University of Pittsburgh, Pittsburgh, PA USA; Department of Pathology, NYU Grossman School of Medicine, New York, NY USA; Department of Medicine, University of Pittsburgh, Pittsburgh, PA USA; UPMC Hillman Cancer Center, Pittsburgh, PA USA; Department of Pathology, Massachusetts General Hospital, Boston, MA USA; Department of Bioengineering, University of Pittsburgh, Pittsburgh, PA USA; Division of Malignant Hematology and Medical Oncology, University of Pittsburgh, Pittsburgh, PA USA

**Author notes:** (Qiangqiang Gu and Riyue Bao co-supervised this work.). Corresponding authors: Qiangqiang Gu;, Riyue Bao. This study was conducted in accordance with protocols approved by the University of Pittsburgh Institutional Review Board.

**Keywords:** Agentic Systems, Nuclear Segmentation, Patient-derived Organoids, Histopathology, Human-In-The-Loop

## Abstract

Nuclear segmentation across patient-derived organoids (PDOs) is complicated by differences in morphology, staining, image scale, and acquisition format. We present ARIPA, an automated human-in-the-loop (HITL) system that coordinates pretrained segmentation tools through a vision-language model (VLM)-based agentic planning, verification, and refinement loop for hematoxylin and eosin images of formalin-fixed, paraffin-embedded PDOs. In contrast to existing tools, ARIPA does not require training and achieves good performance. Tissue-aware tiling and image-derived calibration initialize segmentation; optional expert corrections guide configuration search and light-weight object-level rules without retraining or fine-tuning the pretrained segmentation networks. We evaluated initial ARIPA predictions on more than 50,000 nuclei from 19 images of breast, head and neck, and lung patient-derived cancer PDOs, where nuclear segmentation was challenging due to its 3D structure and variation in imaging layers and malignant cell morphology. Performance was evaluated on agent-selected models before expert feed-back, after feedback from the initial runs excluding direct expert edits, and in comparison to a custom-trained HoVer-Net model previously published by our group. At a minimum polygon inter-section over union (IoU) of 0.5, image-macro F1 was 0.8380 before feedback, 0.9849 after feedback, and 0.4713 for HoVer-Net. Mean area under the F1 score versus IoU curve was 0.8013, 0.9700, and 0.4317, respectively. The mean paired F1 increase after feedback was 0.1468, with increases in 17 out of the total 19 images and no changes in the remaining two. Ground truth nuclei annotations were created by expert review and editing of ARIPA’s post-feed-back outputs. In this cohort, ARIPA outperformed the custom-trained HoVer-Net, and its performance was improved further following expert-guided HITL refinement. By automatic agentic reasoning with VLM review without training, ARIPA has potential to be rapidly deployed across organs and cancers with HITL design, accelerating the fundamental step towards downstream biomedical applications.

## I. Introduction

Patient-derived organoids (PDOs) are ex vivo, three-dimensional (3D) model systems that represent an important class of human-centric New Approach Methodologies (NAMs) for bio-medical research and drug discovery[1]. Unlike traditional cell-based experimental systems, PDOs retain patient-specific molecular and cellular heterogeneity in self-organizing 3D architecture with drug-dependent morphological plasticity[2]. These properties enhance the physiological relevance of PDOs for functional studies of therapeutic response but also introduce substantial variability in their microscopic appearance especially for cancer organoids[3]. Accurate nuclear segmentation is fundamental to artificial intelligence (AI)-driven drug response assessment and the development of functional precision oncology biomarkers at single-cell resolution[3]. Hematoxylin and eosin (H&E) sections preserve cellular morphology and spatial organization of PDOs, but transforming these images into reproducible single-cell measurements remains challenging. Organoid morphology can vary across patients, tumor types, culture conditions, and passages, while their 3D organization can produce substantial differences in nuclear density, sectioning plane, staining appearance, and image quality. Consequently, a segmentation configuration that reliably separates densely packed nuclei in one image may miss pale nuclei, merge adjacent objects, or fragment irregularly stained nuclei in another. Whole slide images (WSIs) additionally capture variation in size, spatial scale, and distribution of PDOs, which are intrinsically inherited from parental tumors.

Nuclear instance segmentation across heterogeneous histopathology images is a longstanding problem in computational pathology[4], [5]. Deep learning approaches can achieve high performance within their training domain, but performance may decrease when biospecimen characteristics differ from those represented during model development[6]. Moreover, tissue-trained models cannot be directly transferred to organoids due to reasons stated above. For laboratories studying PDOs, where manual annotation is generally not available and cohorts may contain samples from multiple tumor types and imaging conditions, repeatedly developing cohort-specific training sets or segmentation models is neither efficient nor scalable.

To address this challenge, we introduce **Agentic Reasoning for Image-based Pathology Analysis (ARIPA)**, a training-free, automated agentic framework for nuclear segmentation that integrates vision-language model (VLM)-guided planning, state-of-the-art segmentation tools, verification and iterative refinement, with optional human-in-the-loop (HITL) feedback. Unlike conventional supervised segmentation workflows, ARIPA implements a stateful workflow in which agents propose segmentation strategies, execute candidate tools, and evaluate the resulting predictions. These evaluations determine whether another refinement cycle is initiated or the automated pass is completed. When expert input is available, corrections are used to guide subsequent configuration choices and light-weight object-level refinement rather than to update model weights. This design is intended to support rapid deployment across heterogeneous biospecimen cohorts without requiring a new labeled dataset for each tissue type, scanner, or imaging condition.

ARIPA’s novelty therefore lies not in introducing a new segmentation architecture, but in shifting adaptation from the model level to the orchestration level. Specifically, ARIPA provides: (i) tissue-aware nuclear segmentation without requiring a cohort-specific annotated training dataset; (ii) agent-guided selection and bounded iterative refinement across segmentation tools and inference configurations; (iii) automated verification using both image-based heuristics and VLM review, and (iv) HITL refinement in which domain-expert edits provide feedback to guide further improvements in nuclear segmentation without model fine-tuning. Tile-level predictions are ultimately assembled into unified slide-level segmentation outputs with seamless integration into downstream digital pathology visualization and analysis in QuPath[7].

We evaluate ARIPA in a cohort of 50,000+ nuclei from 19 H&E images of PDOs originating from lung, head and neck (HN), and breast tumors exhibiting diverse morphologies. Images were acquired using two distinct devices (a Leica Aperio AT2 scanner and a Nikon upright microscope) at 40x magnification. The dataset contains realistic variation in image quality, including regions with reduced focus or local blurring. Performance is assessed before and after expert feedback against ground-truth nuclear annotations for every image, with a separately custom-trained HoVer-Net as a supervised comparator. All tiles from the 19 images were manually inspected to establish the ground truth. This study provides an initial assessment of ARIPA for cross-organ nuclear segmentation, overcoming the traditional barrier of cohort-specific re-training or fine-tuning of its pretrained segmentation networks.

## II. Related work

Most existing nuclear segmentation approaches fall broadly into one of two categories-task-specific models trained on annotated histopathology datasets or broadly pretrained models designed for transfer across imaging domains. Specialized histology models such as HoVer-Net explicitly address nuclei instance segmentation in multi-tissue H&E images[4]. Other approaches have incorporated adversarial learning and synthetic data generation. The conditional generative adversarial network (cGAN)-based multi-organ nuclear segmentation (cGANMNS) was developed using both real and synthetic data[5]. These approaches demonstrate the effectiveness of supervised learning for histologic nuclear segmentation, but their performance remains dependent on the domains represented during model development.

More recent generalist and foundation-model approaches aim to improve transferability across cell types and imaging conditions. Cellpose-Segment Anything Model (Cellpose-SAM) is a pretrained cellular segmentation model developed using large-scale, diverse publicly available datasets[8]. Segment Anything Model (SAM)-based general-purpose approaches provide promptable segmentation capabilities that can be transferred or adapted to biomedical imaging[9], [10]. Recent pathology-specific adaptations, such as PathoSAM, further demonstrate foundation-model approaches for automatic and interactive nuclear instance segmentation[6]. Other architectures, including Common Objects in Context (COCO)-pretrained You Only Look Once (YOLO), provide additional alternatives for object segmentation[11]. These methods vary substantially in their training requirements, prompting strategies, and suitability for nuclear segmentation.

Computational image analysis has also been increasingly applied to PDOs. Prior studies have used fluorescence, brightfield, and other imaging modalities for organoid segmentation, phenotypic characterization, and tumor-versus-normal classification[3], [12], [13], [14]. However, many such approaches rely on application-specific training data or models optimized for a particular imaging modality, tissue type, or experimental setting. This dependency can limit scalability when organoid cohorts span multiple tumor types, culture conditions, imaging platforms, or morphological states.

ARIPA addresses a complementary problem. Rather than introducing another segmentation architecture, it leverages existing segmentation models and conventional image-processing methods as tools within an adaptive reasoning framework. In the current implementation, these include cGANMNS, SAM, YOLO, together with a classical image-processing pipeline based on hematoxylin optical-density extraction, thresholding, morphological operations, and watershed segmentation. ARIPA therefore does not seek to replace specialized histopathology or foundation segmentation models, but that it provides a training-free deployment strategy for coordinating them. By combining heterogeneous segmentation tools, automated verification, iterative refinement, and expert feedback within a stateful agentic workflow, ARIPA is designed to support nuclear segmentation across heterogeneous pathology images without requiring a new cohort-specific annotated training dataset.

## III. Methodology

## A. Task definition and execution graph

ARIPA performs nuclei instance segmentation on histopathology images, identifying individual nuclei and delineating their boundaries. For each tissue-presented image tile extracted from the original image, the system generates an instance label map in which all pixels belonging to a predicted nucleus share a positive integer identifier unique within that tile, while pixels outside the predicted nuclei instances are assigned zero. This representation distinguishes individual nuclei, including adjacent instances. Nuclei boundaries are also converted into polygons and mapped from tile coordinate to the original image coordinates to assemble a slide-level segmentation output.

ARIPA leverages a LangGraph-based[15] workflow to coordinate tissue detection, image tile extraction, planning, nuclear segmentation, and verification. A locally deployed VLM, MedGemma-4B (Google, Mountain View, CA, USA)[16] supports planning, visual assessment, and configuration refinement, while the segmentation tools generate nuclei instance masks. Previously supplied human feedback, when available, informs configuration selection. Nuclear segmentation performance that does not meet the verification threshold triggers configuration refinement and another segmentation iteration, subject to a predefined maximum number of iterations. The highest-scoring segmentation output from each automated refinement cycle undergoes the HITL review. Expert feedback guides configuration adjustments and the learning of rules to suppress false positive detections and recover missed nuclei in subsequent segmentation iterations. This review and refinement cycle continues until the expert requests no further refinement, or the predefined maximum number of feedback iterations is reached. The resulting image tile-level predictions are assembled into slide-level segmentation outputs and summarized in a final report.

**Figure 1.**
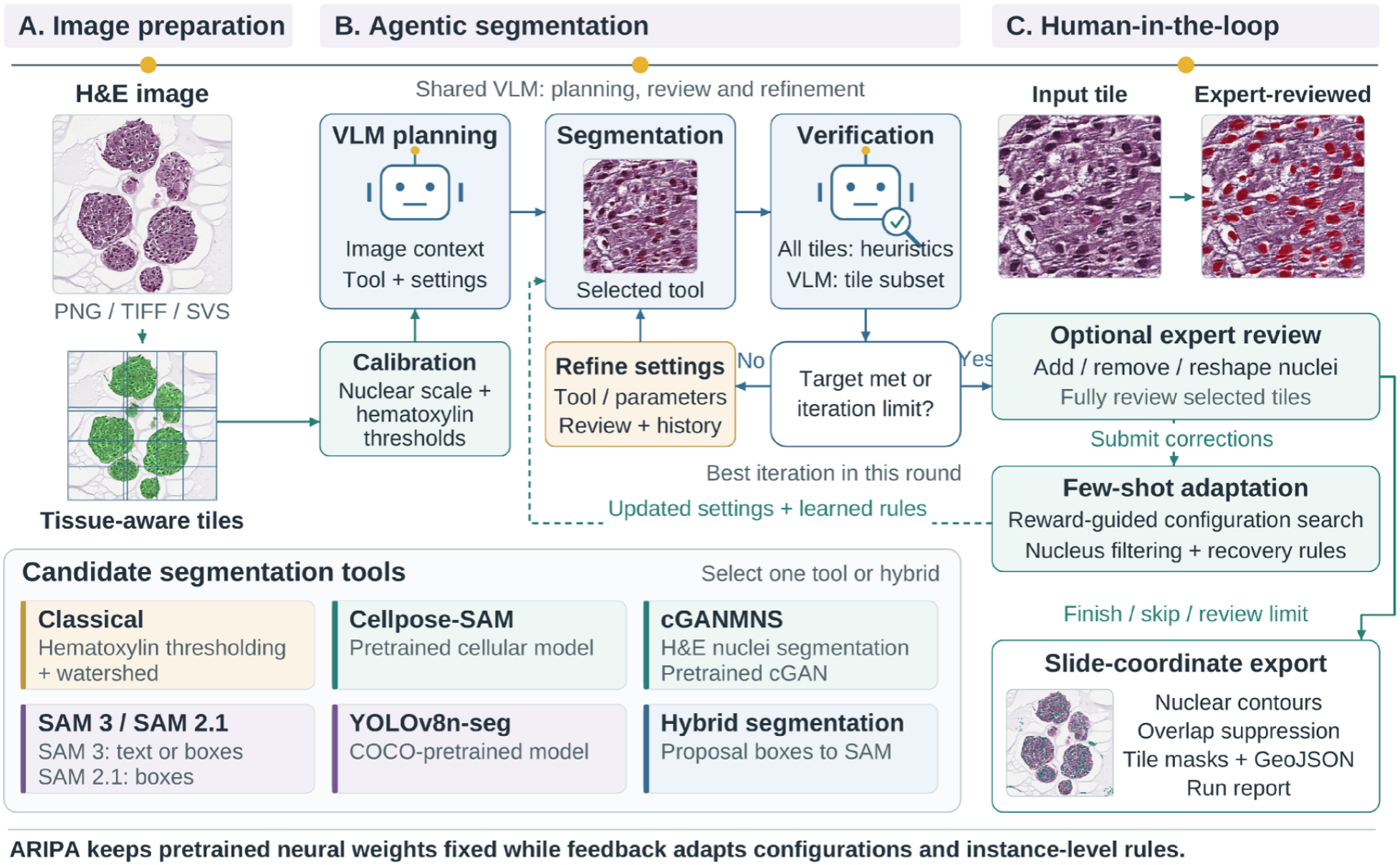
ARIPA workflow integrating tissue-aware pre-processing, segmentation tool selection, automated verification and refinement, HITL adaptation, and slide-coordinate nuclear segmentation export.

### B. Image Pre-processing

Tissue localization identifies the regions to be considered for nuclear segmentation. A binary tissue mask is generated from a thumbnail with a maximum dimension of 2,048 pixels by applying Otsu-based thresholding[17] separately to the saturation channel in Hue, Saturation, Value (HSV) color space and to grayscale intensity. This combination captures both strongly stained tissue and less saturated regions that remain darker than the background. Additional intensity constraints exclude the background non-tissue regions.

To accommodate large tissue areas while retaining local image detail, nuclear segmentation performed on overlapping tiles extracted at native image resolution. The default tile dimensions are up to 1,024 x 1,024 pixels, with a nominal overlap of 64 pixels. A minimum detected tissue fraction of five percent determines tile inclusion, limiting the contribution of background-dominated regions.

Image-derived estimates of staining thresholds and object size guide the initialization of segmentation parameters. These estimates are obtained from up to three tiles with the highest tissue fractions. Within each tile, candidate nuclei regions are identified from the hematoxylin optical-density channel using the upper threshold of three-class multi-Otsu thresholding. Following morphological cleanup, an area-weighted median of connected-component areas provides an estimate of characteristic object size. Median estimates across the sampled tiles inform the initial instance-area limits and watershed marker spacing and constrain subsequent VLM-proposed parameters.

### C. Segmentation tools and dispatch

ARIPA incorporates conventional image processing, pretrained biomedical segmentation models, and general-purpose vision models as candidate segmentation tools. The conventional approach combines thresholding of smoothed hematoxylin optical density with morphological refinement, distance-transform-based watershed segmentation, and instance-area filtering. Cellpose-SAM provides pre-trained biomedical instance segmentation, while cGANMNS employs a pre-trained pix2pix U-Net generator on overlapping 256 x 256-pixel tiles. Its blended contour predictions are converted into nuclei instances through morphological processing and selective watershed separation. The candidate pool also includes SAM-based prompt-guided segmentation and YOLOv8n_seg, a lightweight and efficient general-purpose instance segmentation model pre-trained on COCO dataset[18].

Tool and parameter selection is guided by image context, image-derived calibration, and segmentation quality assessments. The agent could select an individual approach or a hybrid configuration in which bounding boxes derived from initial instance predictions guide SAM-based boundary refinement. Bounding box prompting uses an available SAM3 or SAM 2.1, while text prompting requires SAM3. The selected configuration could be revised during iterative refinement and HITL review. Pre-trained segmentation model weights remain frozen throughout this process, while human feedback informs configuration selection and instance-level filtering and recovery rules. The custom-trained HoVer-Net model is evaluated separately as a supervised comparator.

### D. Verification and iterative refinement

Nuclear segmentation verification combines heuristic assessment of all tile-level predictions with targeted VLM review. The heuristic score considers nuclei size and variability, instance density, mask coverage, and staining characteristics, to assess whether potentially missed nuclei regions remain outside the masks. Under the default configuration, up to six tiles with the highest tissue fractions additionally receive VLM review. For each selected tile, the VLM examines the original tissue image alongside the predicted nuclei masks superimposed, together with numerical segmentation statistics.

At each iteration, the combined verification score for each tile is

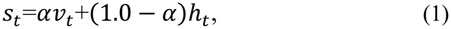

where *ν*_*t*_ and *h*_*t*_ denote the VLM and heuristic scores. Tiles outside the subset selected for VLM review are evaluated using heuristic scores only, giving *S*_*t*_ =*h*_*t*_. The agent score (i.e., *A*_*s*_) is the unweighted mean across tiles. When HITL feedback provides corrected reference masks, the overall score (i.e., *S*) becomes:

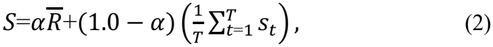

where 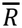 is the aggregated reward measuring agreement with those tasks. Without human feedback, the overall score, *S*, is simply the same as the agent score, *A*_*s*_. Both weighting steps default to α = 0.6, giving greater emphasis to VLM assessment and feedback agreement, respectively. These scores guide refinement and are not calibrated estimates of segmentation accuracy.

Verification findings, image-derived calibration, the current configuration, and iteration history guide subsequent tool selection and parameter adjustment. Each iteration of automated refinement ends when the overall score reaches the acceptance threshold, with 0.80 by default, or the maximum number of iterations is reached. Rewards guide configuration selection during inference, while the pre-trained segmentation models and VLM weights remain frozen.

### E. HITL few-shot adaptation

HITL few-shot adaptation leverages expert-reviewed tiles to guide configuration selection and instance-level refinement. Domain experts add missed nuclei, remove false detections, and adjust nuclei boundaries while retaining correct predictions. The reviewed annotations constitute complete reference masks for their tiles, so the feedback protocol requires full review of each selected tile and treats un-annotated regions as background. Predicted and reference instances are paired using Hungarian assignment algorithm[19] to maximize total IoU, with assigned pairs retained as matches when IoU is greater or equal to 0.5.

For each reviewed tile, the reward (i.e., *R*) is

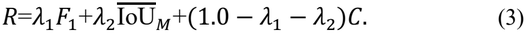

where *F*_1_ measures instance detection agreement, 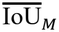 is the mean IoU of matched instances, and *C* measures count agreement. Specifically,

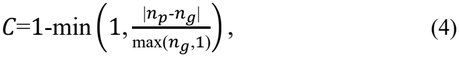

where *n*_*p*_ and *n*_*g*_ denote predicted and reference instance counts. The default weights for λ_1_, and λ_2_ are 0.50, and 0.35, assigning greatest emphasis to detection agreement, followed by spatial overlap, with count consistency providing an additional constraint. The overlap term is zero when no instances are matched, and the reward is set to one when both masks contain no instances. Tile rewards are averaged using weights, max(*n*_*g*_,1) to obtain the aggregate feedback reward, *R*.

Adaptation combines configuration search with example-based filtering and recovery of nuclei instances. Up to eight configurations are evaluated by default, comprising parameter variations and available alternative tools, with an optional additional VLM proposal informed by the feedback. The configuration achieving the highest reward on the reviewed tiles initializes subsequent refinement. Filtering and recovery rules are then learned through nearest-neighbor comparisons of 11 standardized morphological, intensity, and staining features. Additional candidates for missed nuclei are generated by reducing the minimum accepted object area and lowering applicable detection thresholds, such as prediction confidence or hematoxylin intensity thresholds. The candidate masks are then expanded to include neighboring hematoxylin-rich pixels that are not already assigned to predicted nuclei. Rule thresholds are selected jointly according to the feedback reward, and a rule is retained only when it improves agreement on the reviewed tiles. During subsequent segmentation iterations with the same feedback session, enabled rules are applied across all tiles while pre-trained segmentation model weights remain fixed.

### F. Outputs and traceability

ARIPA produces tile-level nuclei instance masks and visualizations, together with geometrically validated nuclei contours exported in GeoJSON using original image coordinates. To limit duplicate detections in overlapping regions, each polygon is retained only when its source tile has the nearest center among tiles containing the centroid of the polygon. This approach consolidates predictions through instance selection without fusing contours across tile boundaries. The refinement history records the segmentation method, parameter settings, and verification results for each iteration, together with expert feedback, allowing changes in the predictions to be traced. Quantitative evaluation leverages agent-generated predictions before and after human feedback, including any applied filtering and recovery rules, while excluding direct substitutions of expert annotations.

## IV. Experimental study

### A. Cohort and evaluation outputs

The study cohort comprises 19 H&E images of formalin-fixed paraffin-embedded (FFPE) PDOs from breast, head and neck, and lung (Table I), each paired with nuclei ground-truth annotations for quantitative evaluations. For each image, initial automated ARIPA predictions, including automated verification and refinement, were compared with post-feedback predictions from the same run. Post-feedback outputs incorporated feedback-driven adaptation but excluded direct substitution of expert-edited annotations. All pre-trained segmentation models included in ARIPA retained their pre-trained weights throughout this study, without cohort-specific fine-tuning.

**TABLE I.** Retained cohort with nuclear ground truth for all 19 images.

| Organ | Input format | Images |
|---|---|---|
| Breast | TIF WSIs | 5 |
| Head and neck | Static Histopathology PNG Images | 5 |
| Head and neck | SVS WSIs | 4 |
| Lung | TIF WSIs | 5 |

### B. Custom-trained HoVer-Net comparator

An independent custom-trained HoVer-Net on head and neck histopathology images from our previous work served as the supervised nuclei segmentation comparator[3]. Its direct predictions, generated without expert feedback or manual editing, were evaluated on all 19 images against the same nuclei reference annotations used for ARIPA evaluations.

The comparison includes three conditions, automated ARIPA before expert feedback, ARIPA after HITL refinement, and direct HoVer-Net predictions, all scored against the same expert-curated ground-truth annotations.

### C. Quantitative evaluation

Nuclear segmentation was evaluated by comparing predicted and reference polygons, with each polygon object representing one nuclei instance. At an IoU threshold of 0.5, maximum-cardinality one-to-one matching identified the largest number of eligible prediction-reference pairs. Matched pairs were counted as true positives (TPs), while unmatched predictions and references were counted as false positives (FPs) and false negatives (FNs), respectively. Precision, recall, F1 score, and instance Jaccard index were defined as

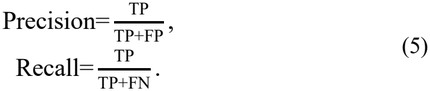

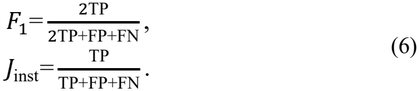

respectively.

To evaluate performance across increasingly stringent spatial-overlap requirements we additionally measured the F1-IoU AUC, where matching was re-computed independently across 11 thresholds within [0.0, 1.0]with increments of 0.1, and F1-IoU AUC was calculated using trapezoidal rule[20], [21]. Positive intersection area was required at every threshold, including zero, preventing disjoint or merely touching polygons from matching. This measure summarizes detection agreement across overlapping thresholds and is distinct from area under the receiver operating characteristic or precision-recall curve.

For each evaluation condition, precision, recall, F1 score, instance Jaccard index, F1-IoU AUC, and absolute nuclei count error, were summarized using unweighted means and sample standard deviations (SDs) across all 19 images and separately by organ. Absolute count error was defined as

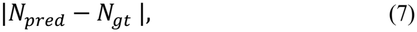

where *N*_*perd*_ and *N*_*gt*_ denote predicted and ground truth reference nuclei counts. Comparisons were paired within each image and conducted separately for each segmentation metric for absolute count error. Feedback-associated differences were calculated by subtracting the pre-feedback from the post-feedback measurements, while comparisons with custom-trained HoVer-Net were calculated by subtracting the HoVer-Net from the ARIPA measurements. Positive differences in nuclear segmentation metrics indicate higher agreement with reference annotations, whereas negative differences in absolute count error indicate improved count agreement. Paired differences were summarized using means and sample SDs, and tested by two-sided Wilcoxon signed-rank test. Pooled precision, recall, F1 score, and instance Jaccard index were calculated from TP, FP, and FN counts summed across images, giving greater influence on images containing more nuclei.

## V. Results

### A. Overall nuclear segmentation performance

Across 19 images containing 54,489 ground truth reference nuclei, F1 score in mean (SD) was 0.8380 (0.2101) for ARIPA before feedback, 0.9849 (0.0194) after feedback, and 0.4713 (0.3485) for custom-trained HoVer-Net. HITL refinement was associated with mean paired increase in mean (SD) of 0.1468 (0.2110) in F1 score, and 0.1687 (0.2194) in F1-IoU AUC (both *P*<0.001, comparing before/after feedback). These comparisons quantify agreement with shared ground truth annotations created through expert review and editing of the post-feedback predictions from ARIPA.

At IoU greater or equal to 0.5, ARIPA before feedback yielded 51,982 TPs, and 2,507 FNs. After feedback, TPs increased to 53,851, while FPs and FNs decreased to 786 and 638, respectively. Custom-trained HoVer-Net yielded 27,139 TPs, 54,219 FPs, and 27,350 FNs. Pooled F1 score was 0.8159 before feedback, 0.9870 after feedback, and 0.3996 for HoVer-Net. Mean absolute count error (SD) was approximately 1,001 (2,217) before feedback, 37 (40) after feedback, and 1,811 (2,772) for HoVer-Net.

Before feedback, ARIPA exceeded HoVer-Net in F1 score on 15 of 19 images (*P* = 0.0053) and in F1-IoU AUC on 16 of 19 images (*P* = 0.0012). After feedback, ARIPA achieved higher precision, recall, F1 score, instance Jaccard index, and F1-IoU AUC, together with lower absolute count error, on all 19 images. Mean (SD) paired differences relative to HoVer-Net were 0.5136 (0.3451) for post-feedback F1 score and 0.5383 (0.2628) for post-feedback F1-IoU AUC (both *P* < 0.001).

### B. Organ-level performance and variability

Organ-level performance results are summarized in Table III for breast, head and neck, and lung images. Mean ARIPA F1 score increased from 0.8869 to 0.9688 in breast, from 0.7697 to 0.9911 in head and neck, and from 0.9121 to 0.9897 in lung, following feedback. Corresponding custom-trained HoVer-Net F1 scores were 0.1698, 0.7423, and 0.2850.

**TABLE II.** Overall nuclear segmentation performance across 19 images, reported as mean (SD).

| Metric | ARIPA before feedback | ARIPA after feedback | Direct HoVer-Net |
|---|---|---|---|
| Precision | 0.7920 (0.2551) | 0.9845 (0.0260) | 0.4508 (0.3458) |
| Recall | 0.9430 (0.0582) | 0.9854 (0.0157) | 0.5149 (0.3475) |
| F1 (IoU threshold 0.5) | 0.8380 (0.2101) | 0.9849 (0.0194) | 0.4713 (0.3485) |
| Instance Jaccard | 0.7636 (0.2509) | 0.9708 (0.0357) | 0.3774 (0.3226) |
| F1-IoU AUC [0,1] | 0.8013 (0.2149) | 0.9700 (0.0242) | 0.4317 (0.2645) |
| Absolute count error (nuclei) | 1000.53 (2216.71) | 37.05 (39.56) | 1811.42 (2771.62) |

**TABLE III.** Organ-level nuclear segmentation performance of ARIPA before and after expert feedback and HoVer-net, reported as mean (SD).

| Organ (images), stage/method | Precision | Recall | F1 | Instance Jaccard | F1-IoU AUC |
|---|---|---|---|---|---|
| Breast (5), before | 0.8476 (0.1563) | 0.9463 (0.0509) | 0.8869 (0.0965) | 0.8074 (0.1521) | 0.8733 (0.0952) |
| Breast (5), after | 0.9670 (0.0435) | 0.9708 (0.0225) | 0.9688 (0.0324) | 0.9409 (0.0586) | 0.9548 (0.0364) |
| Breast (5), HoVer-Net | 0.1244 (0.0469) | 0.2742 (0.1292) | 0.1698 (0.0680) | 0.0940 (0.0421) | 0.2197 (0.0442) |
| Head and neck (9), before | 0.7106 (0.3424) | 0.9399 (0.0662) | 0.7697 (0.2877) | 0.6955 (0.3388) | 0.7361 (0.2921) |
| Head and neck (9), after | 0.9947 (0.0081) | 0.9876 (0.0095) | 0.9911 (0.0064) | 0.9825 (0.0126) | 0.9720 (0.0174) |
| Head and neck (9), HoVer-Net | 0.7260 (0.2768) | 0.7641 (0.2892) | 0.7423 (0.2798) | 0.6404 (0.2570) | 0.6282 (0.2215) |
| Lung (5), before | 0.8830 (0.0700) | 0.9452 (0.0620) | 0.9121 (0.0589) | 0.8426 (0.0990) | 0.8466 (0.0982) |
| Lung (5), after | 0.9835 (0.0201) | 0.9961 (0.0020) | 0.9897 (0.0107) | 0.9798 (0.0209) | 0.9816 (0.0157) |
| Lung (5), HoVer-Net | 0.2820 (0.2090) | 0.3069 (0.3053) | 0.2850 (0.2511) | 0.1875 (0.1830) | 0.2902 (0.2162) |

Across individual images, ARIPA F1 score ranged from 0.2447 to 0.9984 before feedback and from 0.9115 to 0.9995 after feedback. Custom-trained HoVer-Net F1 score ranged from 0.0007 to 0.9416. Minimum F1-IoU AUC values were 0.2176 before feedback, 0.8905 after feedback, and 0.0120 for HoVer-Net. The largest absolute nuclei count errors were 8,799 for initial ARIPA predictions, 159 for post-feedback predictions, and 9,553 for HoVer-Net. These results indicate more consistent agreement with the reference annotations across the evaluated images following HITL refinement.

### C. Changes following HITL refinement

Following human feedback, precision, F1, instance Jaccard index, and F1-IoU AUC increased in 17 images and remained unchanged in HN3 and HN9. Recall increased in 14 images, remained unchanged in four, and decreased slightly for BC5 (0.9889 to 0.9875). Absolute nuclei count error decreased in 16 images, remained unchanged in two, and increased in BC4 (44 to 90 nuclei). The largest F1 score improvements occurred in HN8 (0.2447 to 0.9802) and HN6 (0.3730 to 0.9985), accompanied by reductions in FPs from 9,109 to 48 and from 5,303 to 1, respectively. These comparisons evaluated ARIPA predictions and excluded direct substitution of expert-edited annotations. However, the reference annotations were derived from expert review and editing of the post-feedback predictions from ARIPA. The observed improvements therefore describe changes within the evaluated runs without independently establishing the causal benefit of human feedback or its transfer to unedited regions.

## VI. Discussion and limitations

ARIPA enables nuclei segmentation without cohort-specific training of segmentation backbones or the development of a new segmentation architecture. The framework integrates conventional image processing and pre-trained general-purpose or biomedical specialized models through agentic planning, verification, and refinement. Across breast, head and neck, and lung images, initial ARIPA predictions achieved a mean F1 score of 0.8380, compared with 0.4713 for custom-trained HoVer-Net predictions. Following HITL refinement, mean F1 score increased to 0.9849, with improvements in 17 images and unchanged performance in the remaining two images. Expert feedback guided configuration selection and instance-level filtering and recovery while pre-trained network weights remained fixed. These findings support the applicability of ARIPA across the evaluated organ groups and demonstrate improved agreement with ground truth annotations.

Ground truth annotations were generated through the interactive user interface of ARIPA, where experts added missed nuclei, removed false detections, and corrected the boundaries of post-feedback predictions. This annotation strategy was adopted since completely independent manual delineation of approximately 55,000 nuclei across all 19 images was time-consuming and impractical. Nevertheless, editing existing predictions rather than independently delineating nuclei may introduce annotation bias, which future study plans to assess through evaluation against independently generated reference annotations. Given such, the near-ceiling post-feedback agreement is expected in part by construction; pre-feedback and organ-level trends better reflect the agentic framework’s standalone contribution. In addition, considering HoVer-Net was trained on head and neck data and applied to breast and lung without retraining, its lower performance there partly reflects out-of-domain transfer rather than a general supervised-learning limitation; we treat this as illustrating the retraining burden of cohort-specific supervised models, not a controlled benchmark against agentic methods. Lastly, the 19-image cohort reflects the complexity, time, and expense of generating multi-cancer PDO histological data, and this study serves as proof-of-concept for training-free, scalable agentic reasoning on difficult-to-segment tumor PDO images at single cell resolution.

## VII. Conclusion

ARIPA introduces a multi-agentic framework for nuclear segmentation that integrates existing segmentation tools with automated planning, verification, iterative refinement, and HITL adaptation. Its principal contribution is enabling image-specific tool selection and refinement without developing a new segmentation architecture or fine-tuning pre-trained segmentation backbones on the target dataset. Expert feedback guides configuration selection on instance-level refinement, improving agreement with reference annotations while preserving pretrained network weights. Evaluation across breast, head and neck, and lung organoid histology supports the feasibility of this approach across multiple organ systems and its potential to reduce reliance on customized model training. Future work will further delineate the contributions of tool selection, iterative refinement, and instance-level adaptation; assess sensitivity to image scale and verification settings; and quantify the expert effort required to achieve the observed performance. Expanded cohorts, greater tissue diversity, and independently generated annotations will further evaluate generalizability and determine the benefits of HITL refinement.

## Acknowledgment

R.B. conceived the biological question motivating the study, provided data, and co-supervised the study with Q.G. Q.G. designed, implemented, and validated the agentic framework, and performed the experimental analysis. R.D. developed head and neck cancer and lung cancer PDOs. R.D. also performed sample processing and captured images for all cases in this study. J.C., A.S., B.I., and I.K.Z. assisted with the testing of the frame-work. R.X., S.J.S., and A.D.S. contributed pathological insights. Q.G. drafted the manuscript, and R.B. reviewed and edited it. All authors read and approved the final manuscript.

The authors thank the clinical research teams of the University of Pittsburgh Head and Neck Cancer SPORE program and lung cancer surgical program for facilitating patient consent and tumor specimen collection. The authors acknowledge the University of Pittsburgh organoid core (Director: Dr. Polly E. Mattila) for providing the breast cancer PDOs, and thank Dr. Adrian Lee and his laboratory for sharing research resources.

## References

[1] W. Liu, P. D. Pang, C. A. Wu, D. Tagle, and J. C. Wu, “New approach methodologies for drug discovery,” Cell, vol. 189, no. 7, pp. 1877–1903, Apr. 2026, doi: 10.1016/j.cell.2026.02.012.

[2] J. Drost and H. Clevers, “Organoids in cancer research,” Nat. Rev. Cancer, vol. 18, no. 7, pp. 407–418, Jul. 2018, doi: 10.1038/s41568-018-0007-6.

[3] R. Doerfler et al., “Integrating artificial intelligence-driven digital pathology and genomics to establish patient-derived organoids as new approach methodologies for drug response in head and neck cancer,” Oral Oncol., vol. 171, p. 107742, Dec. 2025, doi: 10.1016/j.oraloncology.2025.107742.

[4] S. Graham et al., “Hover-Net: Simultaneous segmentation and classification of nuclei in multi-tissue histology images,” Med. Image Anal., vol. 58, p. 101563, Dec. 2019, doi: 10.1016/j.media.2019.101563.

[5] F. Mahmood et al., “Deep Adversarial Training for Multi-Organ Nuclei Segmentation in Histopathology Images,” IEEE Trans. Med. Imaging, vol. 39, no. 11, pp. 3257–3267, Nov. 2020, doi: 10.1109/TMI.2019.2927182.

[6] T. Griebel, A. Archit, and C. Pape, “Segment Anything for Histopathology,” in Proceedings of The 8th International Conference on Medical Imaging with Deep Learning, PMLR, May 2026, pp. 515–546. Accessed: Sep. 26, 2026. [Online]. Available: https://proceedings.mlr.press/v301/griebel26a.html

[7] P. Bankhead et al., “QuPath: Open source software for digital pathology image analysis,” Sci. Rep., vol. 7, p. 16878, Dec. 2017, doi: 10.1038/s41598-017-17204-5.

[8] M. Pachitariu, M. Rariden, and C. Stringer, “Cellpose-SAM: superhuman generalization for cellular segmentation,” May 01, 2025, bioRxiv. doi: 10.1101/2025.04.28.651001.

[9] N. Ravi et al., “SAM 2: Segment Anything in Images and Videos,” presented at the The Thirteenth International Conference on Learning Representations, Oct. 2024. Accessed: Sep. 26, 2026. [Online]. Available: https://openreview.net/forum?id=Ha6RTeWMd0

[10] N. Carion et al., “SAM 3: Segment Anything with Concepts,” Mar. 28, 2026, arXiv: arXiv:2511.16719. doi: 10.48550/arXiv.2511.16719.

[11] R. Varghese and S. M., “YOLOv8: A Novel Object Detection Algorithm with Enhanced Performance and Robustness,” in 2024 International Conference on Advances in Data Engineering and Intelligent Computing Systems (ADICS), Apr. 2024, pp. 1–6. doi: 10.1109/ADICS58448.2024.10533619.

[12] P. Mukashyaka et al., “High-throughput deconvolution of 3D organoid dynamics at cellular resolution for cancer pharmacology with Cellos,” Nat. Commun., vol. 14, no. 1, p. 8406, Dec. 2023, doi: 10.1038/s41467-023-44162-6.

[13] H. T. Ong et al., “Digitalized organoids: integrated pipeline for high-speed 3D analysis of organoid structures using multilevel segmentation and cellular topology,” Nat. Methods, vol. 22, no. 6, pp. 1343–1354, Jun. 2025, doi: 10.1038/s41592-025-02685-4.

[14] T. Park, T. K. Kim, Y. D. Han, K.-A. Kim, H. Kim, and H. S. Kim, “Development of a deep learning based image processing tool for enhanced organoid analysis,” Sci. Rep., vol. 13, no. 1, p. 19841, Nov. 2023, doi: 10.1038/s41598-023-46485-2.

[15] D. Pearson, S. Shapiro, E. S. G. Venegas, S. Al-Khatib, and A. P. Arzola, “Graph-Based Agentic AI with LangGraph: Workflow Pathways for Long-Running Stateful Business Processes,” Jul. 21, 2026, arXiv: arXiv:2607.19297. doi: 10.48550/arXiv.2607.19297.

[16] A. Sellergren et al., “MedGemma Technical Report,” Apr. 06, 2026, arXiv: arXiv:2507.05201. doi: 10.48550/arXiv.2507.05201.

[17] N. Otsu, “A Threshold Selection Method from Gray-Level Histograms,” IEEE Trans. Syst. Man Cybern., vol. 9, no. 1, pp. 62–66, Jan. 1979, doi: 10.1109/TSMC.1979.4310076.

[18] D. Reis, J. Kupec, J. Hong, and A. Daoudi, “Real-Time Flying Object Detection with YOLOv8,” May 22, 2024, arXiv: arXiv:2305.09972. doi: 10.48550/arXiv.2305.09972.

[19] H. W. Kuhn, “The Hungarian method for the assignment problem,” Nav. Res. Logist. Q., vol. 2, no. 1–2, pp. 83–97, 1955, doi: 10.1002/nav.3800020109.

[20] R. J. Tallarida and R. B. Murray, “Area under a Curve: Trapezoidal and Simpson’s Rules,” in Manual of Pharmacologic Calculations: With Computer Programs, R. J. Tallarida and R. B. Murray, Eds., New York, NY: Springer, 1987, pp. 77–81. doi: 10.1007/978-1-4612-4974-0_26.

[21] I. Safder et al., “Parsing AUC Result-Figures in Machine Learning Specific Scholarly Documents for Semantically-enriched Summarization,” Appl. Artif. Intell., vol. 36, no. 1, p. 2004347, Dec. 2022, doi: 10.1080/08839514.2021.2004347.

